# Urine Proteomics in Methamphetamine Addicted Patients

**DOI:** 10.1101/2024.06.18.599273

**Authors:** Ziyun Shen, Juncheng Liang, Shuxuan Tang, Xuanzhen Pan, Yijin Bao, Yanping Deng, Youhe Gao

**Author notes:** Fund Project: Beijing Normal University (11100704). Shen and Liang contributed equally to this work. Corresponding author: Gao Youhe, male, professor. Yanping Deng, female, professor.

## Abstract

Drug addiction is a serious chronic relapsing brain disease, and methamphetamine dependence has a complex course and is difficult to treat, causing a serious public health burden. In this study, we compared and analyzed the urine proteome of methamphetamine-using patients, methamphetamine-withdrawing patients, healthy individuals. The urine proteome of methamphetamine-using patients and methamphetamine-withdrawing patients was significantly different from that of healthy individuals, and some differential proteins and their enriched biological functions showed that they were associated with addiction or neurotoxicity of methamphetamine. The fact, that the urine proteome of patients who withdrew methamphetamine for more than 3 months were significantly different from that of healthy patients, may reveal the reasons for the high rate of methamphetamine relapse and potential intervention targets. This study proved that urine proteome can reflect the effects of addiction on the body comprehensively, and has the potential to provide clues for addiction.

## 1 Introduction

Drug addiction is a global, major public health and social problem. According to the World Drug Report 2021, in 2020, approximately 284 million people aged 15-64 worldwide will be using drugs, an increase of 22 percent over the past decade. While drug addiction seriously jeopardizes the physical and mental health of drug users, the continued growth of the addicted population places a huge health, economic and security burden on society. At present, the International Convention on Narcotics Control divides addictive drugs into three categories: (1) narcotic drugs, including opioids, cocaine and cannabis; (2) psychotropic drugs, including sedative-hypnotics, central stimulants and hallucinogens; and (3) other drugs, including tobacco, alcohol and volatile organic solvents[1].

Methamphetamine (METH) is a strongly addictive drug that has caused serious social and health problems worldwide. In China, there are nearly 1.5 million drug addicts, including nearly 800,000 methamphetamine abusers, and methamphetamine has replaced heroin as the most abused drug in China. Methamphetamine has replaced heroin as the most abused drug in China[2]. Long-term abuse of methamphetamine can cause serious physical and mental harm. Methamphetamine is highly lipophilic[3]. Compared to other psychostimulants, methamphetamine enters the central nervous system (CNS) more easily and is distributed throughout the brain for a longer duration of action, making methamphetamine more addictive[4]. Methamphetamine induces long-term neurotoxicity with severe and long-lasting damage to the CNS[5]. Methamphetamine induces long-term neurotoxicity, which causes severe and lasting damage to the central nervous system. Methamphetamine addicts often suffer long-term psychiatric disorders and cognitive decline[6]. The neurochemical mechanisms of neurotoxicity and excitotoxicity are complex and unknown.

Research has shown that methamphetamine addiction involves changes in multiple neurotransmitter systems and neural circuits. The chemical structure of methamphetamine is similar to that of monoamines[7]. In the brain, methamphetamine acts on neurotransmitter systems such as dopamine, 5-hydroxytryptamine, norepinephrine, and epinephrine, leading to alterations in neurotransmitter release and reuptake, which produces intense feelings of euphoria and addictive behavior[8]. The final result of methamphetamine’s effects is a strong sense of euphoria and addictive behavior. The end result of methamphetamine’s effects is the overstimulation of monoaminergic pathways in the central and peripheral nervous systems, leading to severe dysfunction and even neuronal degeneration in multiple brain regions such as the striatum, prefrontal cortex, and hippocampus, which further exacerbates the development and maintenance of addictive behaviors[7]. The formation and maintenance of addictive behaviors are further exacerbated. Methamphetamine dependence is a complex and difficult disease to treat, resulting in a significant public health burden[9]. The disease is complex and difficult to treat, resulting in a serious public health burden.

Drug addiction is a serious chronic relapsing brain disease involving higher neural activities such as reward, motivation, learning memory and decision-making, and its mechanism is very complex, involving the participation of multiple brain regions, circuits and systems. The mechanism of drug addiction has not yet been fully elucidated, which makes the prevention and treatment of addiction more difficult. Therefore, understanding the effects of drug addiction on the organism and searching for related biomarkers are of great significance for the development of effective intervention and treatment strategies.

Drug addiction is a dynamic behavioral and physiological process. Metabolomics technology and proteomics technology are widely used in the study of addictive substances. Proteomics is a global approach to study the expression and function of proteins in living organisms, and the use of proteomics technology currently enables the analysis of protein expression and alterations during drug addiction. Proteomics techniques have been widely used to study the effects of a wide range of addictive substances on animal models and humans; for example, researchers have used proteomics techniques to analyze the plasma of patients addicted to methamphetamine[10], or to assess protein expression in the brain tissue of animals and humans exposed to addictive substances such as alcohol, amphetamines, methamphetamine, cocaine, marijuana, morphine, nicotine, etc.[11] These studies provide a rich resource for further research on addiction-related biochemical pathways, genes, and proteins. However, few researchers have used urine proteomics to study drug addiction so far.

Proteins in urine contain a wealth of information that can reflect small changes produced in different systems and organs of the organism. Compared with other body fluid samples, urine is not affected by homeostatic mechanisms and is able to accumulate early changes in the physiological state of the organism, which is highly sensitive and indicative, and has the potential to assist in the early diagnosis, treatment and prognostic monitoring of diseases. In addition, thanks to the non-invasive collection method, urine can be collected continuously, in large quantities, repeatedly and stored conveniently and stably, and the components are relatively simple and easy to analyze. This makes urine an ideal sample for the detection of biomarkers of disease[12]. The results of this study are summarized in the following table.

Numerous studies of animal models and clinical samples illustrate that urine proteomics studies can provide clues for early diagnosis and intervention in brain disorders such as neurodegenerative diseases[13]. Zhang et al. characterized the urinary proteome of mice with Alzheimer’s disease and found that Alzheimer’s disease-related biomarkers were present in urine before pathology showed β-amyloid deposition[14]. Song et al. found that urinary exosomal proteins in a 5XFAD mouse model exhibited Alzheimer’s disease-related differences before amyloid plaque deposition was detected[15]. Matthias Mann’s research team combined proteomics technology, genetic screening, and machine learning, and the urine proteome can distinguish people who carry different Parkinson’s disease-associated mutation genes with different disease manifestations, and can be applied to stratify familial Parkinson’s disease[16].

In addition, urine proteomics has been studied in mental disorders and behavioral disorders, etc. Meng et al. found that a screened urine protein biomarker panel could effectively differentiate between healthy and autistic children of different age groups, which has the potential to assist in the early diagnosis and intervention of autism, and validated the results using a randomized group approach[17]. Wang et al. found that the proteome of autistic patients showed changes in biological pathways related to disease mechanisms[18]. Huan et al. detected differentially expressed proteins in patients with major depressive disorder with different responses to antidepressant medications, and urinary biomarkers have the potential to predict effective therapeutic interventions for patients with major depressive disorder, providing clues and rationale for precision treatment and improving patients’ quality of life[19].

No urine proteomic findings involving addictive drugs such as methamphetamine have been published. Given that the urine proteome can reflect changes resulting from a wide range of brain diseases, this study hopes to broaden the potential application of the urine proteome to drug addiction, a complex and intractable brain disease.

The aim of this study is to investigate the differences between the urinary proteome of methamphetamine-addicted patients and that of healthy people, and to systematically analyze the differences at different levels, such as biological pathways and behavioral characteristics of the population, with the aim of revealing the effects of methamphetamine addiction on the physiological state of the body, and to search for potential protein biomarkers related to methamphetamine addiction and candidate drug targets for the treatment of methamphetamine addiction, thereby providing new clues for the diagnosis and treatment of drug addiction, and providing a new perspective for the study of drug addiction. treatment of methamphetamine addiction, thus providing new clues for the diagnosis and treatment of methamphetamine addiction, as well as providing a new perspective for the study of drug addiction.

## 2 Materials and Methods

### 2.1 Experimental materials

#### 2.1.1 Experimental consumables

1.5 ml/2 ml centrifuge tube (Axygen, USA), 50 ml/15 ml centrifuge tube (Corning, USA), 96-well cell culture plate (Corning, USA), 10 kD filter (Pall, USA), Oasis HLB solid-phase extraction column (Waters, USA), 1 ml/200 ul/20 ul pipette tips (Axygen, USA), BCA kit (Thermo Fisher Scientific, USA), high pH reverse peptide isolation kit (Thermo Fisher Scientific, USA), iRT (indexed retention time, BioGnosis, UK).

#### 2.1.2 Experimental apparatus

Freezing high-speed centrifuge (Thermo Fisher Scientific, USA), vacuum concentrator (Thermo Fisher Scientific, USA), DK-S22 electric thermostatic water bath (Shanghai Jinghong Experimental Equipment Co., Ltd.), full-wavelength multifunctional enzyme labeling instrument (BMG Labtech, Germany), oscillator (Thermo Fisher Scientific, USA), TS100 constant temperature mixer (Hangzhou Ruicheng Instrument Co. Thermo Fisher Scientific), TS100 constant temperature mixer (Hangzhou Ruicheng Instrument Co., Ltd.), electronic balance (METTLER TOLEDO, Switzerland), −80 ultra-low-temperature freezer refrigerator (Thermo Fisher Scientific, U.S.A.), EASY-nLC1200 ultra-high performance liquid Chromatography (Thermo Fisher Scientific, USA), Orbitrap Fusion Lumos Tribird Mass Spectrometer (Thermo Fisher Scientific, USA).

#### 2.1.3 Experimental reagents

Trypsin Golden (Promega, USA), dithiothreitol DTT (Sigma, Germany), iodoacetamide IAA (Sigma, Germany), pure water (Wahaha, China), mass spectrometry grade methanol (Thermo Fisher Scientific, USA), mass spectrometry grade acetonitrile (Thermo Fisher Scientific, USA), Tris-Base (Thermo Fisher Scientific, USA) and other reagents were used. Thermo Fisher Scientific), mass spectrometry grade purified water (Thermo Fisher Scientific, USA), Tris-Base (Promega, USA) and other reagents.

#### 2.1.4 Analysis software

Proteome Discoverer (Version2.1, Thermo Fisher Scientific, USA), Spectronaut Pulsar (Biognosys, UK), Ingenuity Pathway Analysis (Qiagen, Germany); R studio (Version1.2.5001); Xftp 7; Xshell 7.

### 2.2 Experimental Methods

#### 2.2.1 Study population selection and design

Subjects were divided into 3 groups: the first group was the acute group, i.e., patients who stopped using methamphetamine acutely for less than 24 hours and were admitted to the hospital for treatment, denoted as A; the second group was the recovery group, i.e., patients who stopped using drugs for more than 3 months and were admitted to the recovery treatment, denoted as R; and the third group was the control group, i.e., healthy volunteers, denoted as H. In collaboration with hospitals and research institutes, the urine samples were collected, and the collected urine samples were were temporarily stored in a −80°C refrigerator for backup. A total of 22 urine samples from patients in the acute stage, 26 samples from patients in the recovery stage, and 28 urine samples from healthy people were collected. Some of the samples were excluded because of their low peptide content or because physical examination showed that they had other diseases. Finally, 19 urine samples from patients in the acute stage, 22 urine samples from patients in the rehabilitation stage, and 25 urine samples from healthy people were included in the analysis. The study was reviewed and approved by the Ethics Committee of Peking University School of Medicine, and all subjects signed an informed consent form.

**Fig. 1.**
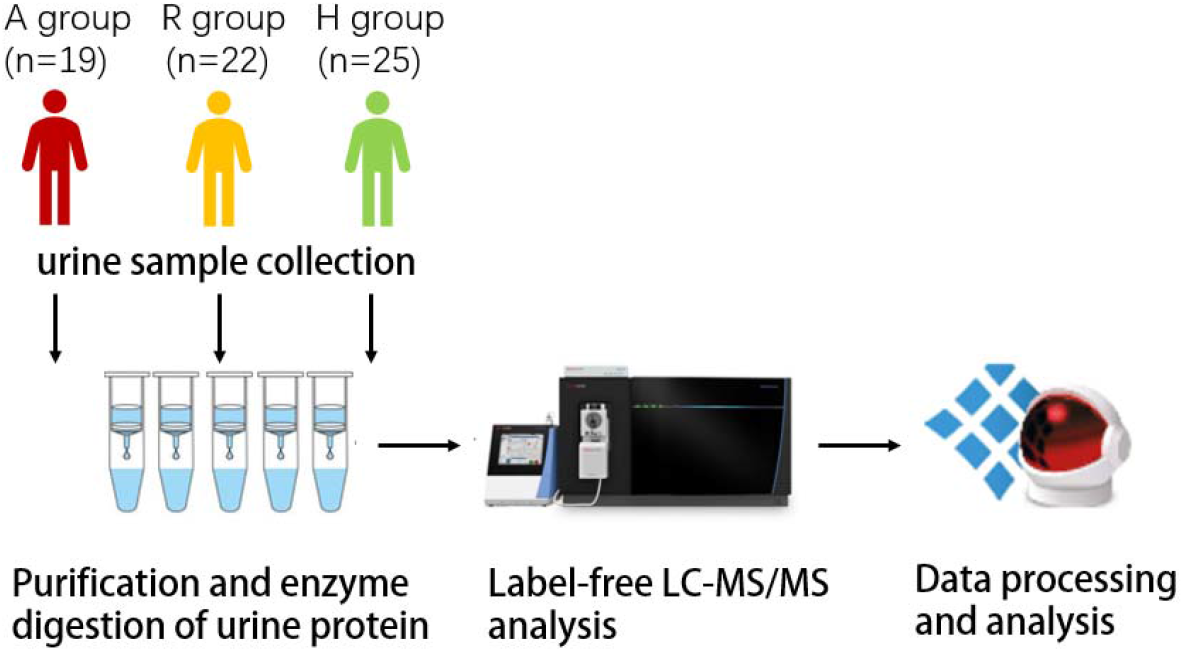
Research methodology and technical route

#### 2.2.2 Urine sample processing

Remove 2ml of urine samples and thaw, centrifuge at 4, 12000×g for 30 minutes, remove cellular debris, take the supernatant and add 1M dithiothreitol (DTT, Sigma) storage solution 40ul, to reach the working concentration of DTT 20mM, mix well and then heated in a metal bath at 37 for 60 minutes, cool to room temperature, add iodoacetamide (Iodoacetamide, IAA, Sigma) storage solution 100ul, to reach the working concentration of IAM, mix well and then react for 45 minutes at room temperature and protected from light. Iodoacetamide, IAA, Sigma) reservoir solution 100ul, to reach the working concentration of IAM, mix well, and then the reaction was carried out for 45 minutes at room temperature and protected from light. At the end of the reaction, the samples were transferred to new centrifuge tubes, mixed thoroughly with three times the volume of pre-cooled anhydrous ethanol, and placed in a refrigerator at −20°C for 24 h to precipitate the proteins. At the end of precipitation, centrifuge the sample at 4 for 30 minutes at 10,000×g, discard the supernatant, dry the protein precipitate, and add 200ul of 20mM Tris solution to the protein precipitate to reconstitute it. After centrifugation, the supernatant was retained and the protein concentration was determined by the Bradford method. Using the filter-assisted sample preparation (FASP) method, urinary protein extracts were added to the filter membrane of a 10kD ultrafiltration tube (Pall, Port Washington, NY, USA), washed three times with 20mM Tris solution, respectively, and the protein was re-solubilized by the addition of 30mM Tris solution, and the protein was added in a proportional manner (urinary protein: trypsin = 50:1) to each sample. Trypsin (Trypsin Gold, Mass Spec Grade, Promega, Fitchburg, WI, USA) was digested and incubated at 37°C for 16 h. The digested filtrate was the peptide mixture. The collected peptide mixture was desalted by an Oasis HLB solid phase extraction column and dried under vacuum, and stored at −80°C. The peptide mixture was then extracted from the peptide mixture with a 0.5 μl of 0.1 % PBDE. The lyophilized peptide powder was re-dissolved by adding 30 μL of 0.1% formic acid water, and then the peptide concentration was determined by using the BCA kit, and the peptide concentration was diluted to 0.5 μg/μL, and 4 μL of each sample was taken out as the mix sample.

#### 2.2.3 LC-MS/MS tandem mass spectrometry analysis

All identification samples were added to a 100-fold dilution of iRT standard solution at a volume ratio of 20:1 sample:iRT, and retention times were standardized. Data Independence Acquisition (DIA) was performed on all samples, and each sample was repeated 2 times, with 1-mix samples inserted every 10 pins as a quality control. The 1ug samples were separated using EASY-nLC1200 liquid chromatography (elution time: 90min, gradient: mobile phase A: 0.1% formic acid, mobile phase B: 80% acetonitrile), and the eluted peptides were entered into the Orbitrap Fusion Lumos Tribird mass spectrometer for analysis, and the corresponding raw files of the samples were generated.

#### 2.2.4 Data processing and analysis

The raw files collected in DIA mode were imported into Spectronaut software for analysis, and the highly reliable protein standard was peptide q value<0.01. The peak area quantification method was applied to quantify the protein by applying the peak area of all fragmented ion peaks of secondary peptides, and the automatic normalization was processed.

Proteins containing two or more specific peptides were retained, and the missing values were replaced with 0. The content of different proteins identified in each sample was calculated, and the different samples were compared to screen for differential proteins according to the screening conditions of FC ≥ 1.5 & ≤ 0.67, P < 0.05.

Unsupervised cluster analysis (HCA), principal component analysis (PCA), and OPLS-DA analysis were performed using the Wukong platform (https://omicsolution.org/wkomics/main/). Functional enrichment analysis of differential proteins was performed using the DAVID database (https://david.ncifcrf.gov/), and the results of 3 aspects of biological process, cellular localization and molecular function were obtained. And the differential proteins were analyzed by Ingenuity Pathway Analysis (IPA). The search of differential proteins and related pathways was performed based on Pubmed database (https://pubmed.ncbi.nlm.nih.gov/). Protein Interaction Network Analysis (PIN) was performed using STRING database (https://cn.string-db.org/).

## 3 Results and Discussion

### 3.1 Basic information about the sample

We identified a total of 2917 highly plausible urinary proteins (unique peptides ≥2, FDR <1%) by mass spectrometry using a non-labeled quantitative technique in a data-independent acquisition mode in 19 acute-phase patient samples, 22 convalescent-phase patient samples, and 25 healthy volunteer samples.

**Table 1.** Identification of samples.

| Class | A group | R group | H group | Total |
| --- | --- | --- | --- | --- |
| Number of samples | 19 | 22 | 25 | 66 |
| Age (Years) | 34±5 | 33±7 | 31±5 | 32±6 |
| Average of protein groups | 2057±335 | 2013±268 | 2052±303 | 2040±302 |

### 3.2 Comparative analysis of acute and healthy groups

#### 3.2.1 Differential protein analysis

In the comparative analysis between the healthy group and the group of patients in the acute phase, 143 differential proteins were screened for FC >1.5 or <0.67, P < 0.05.

Six of the differential protein FCs were 0, showing a variation from present to absent, i.e., a value of 0 was detected in the urine of all methamphetamine-addicted patients, and was detected in the urine of several healthy individuals.

**Figure 2.**
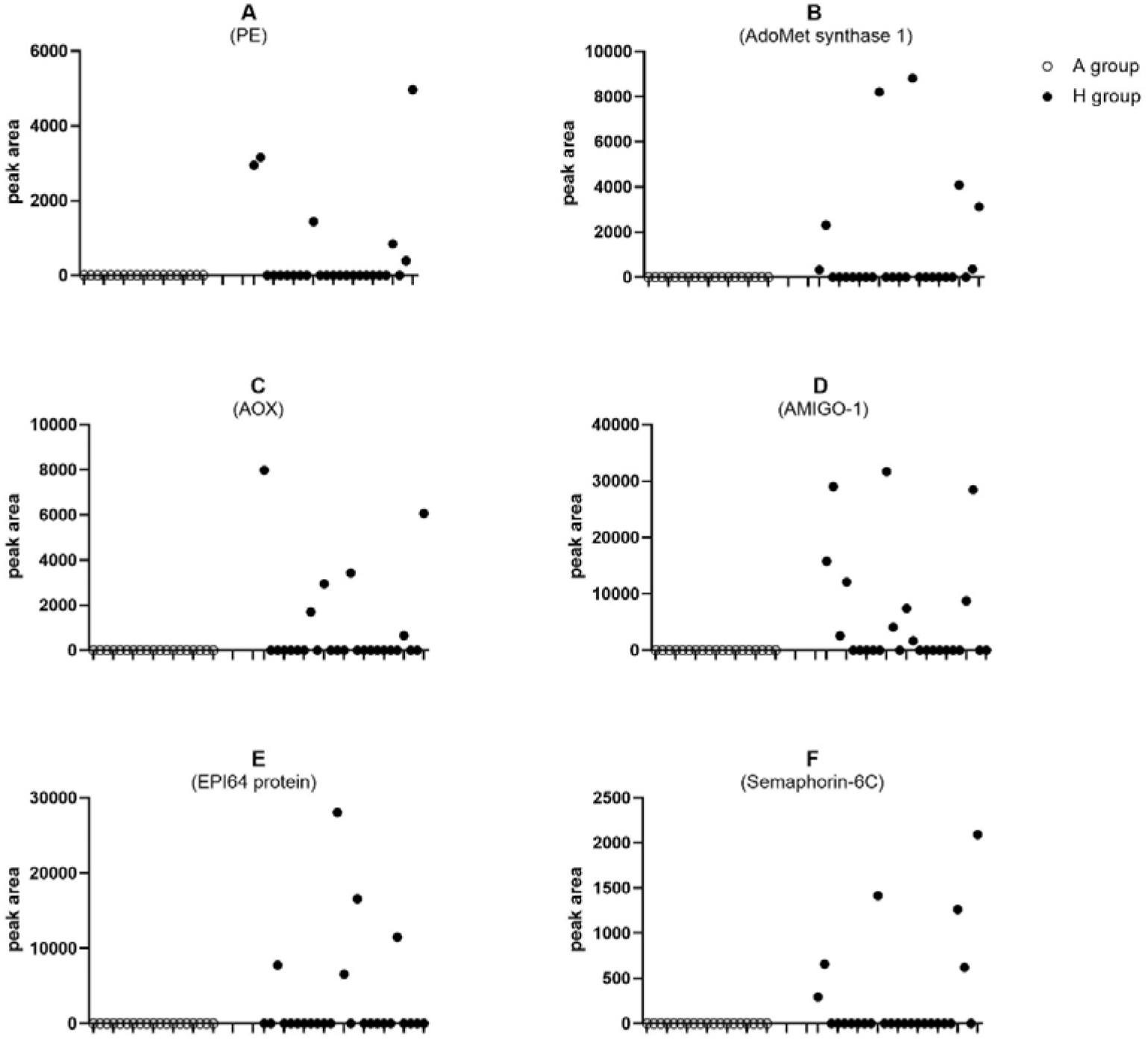
Proteins with a multiplicity of change (FC) of 0 in the comparison of methamphetamine patients and healthy individuals (p < 0.05)

#### 3.2.2 IPA pathway analysis of differential proteins

Differential proteins were enriched to 38 IPA pathways, many of which are closely associated with mental illness and drug addiction.

**Table 2.**
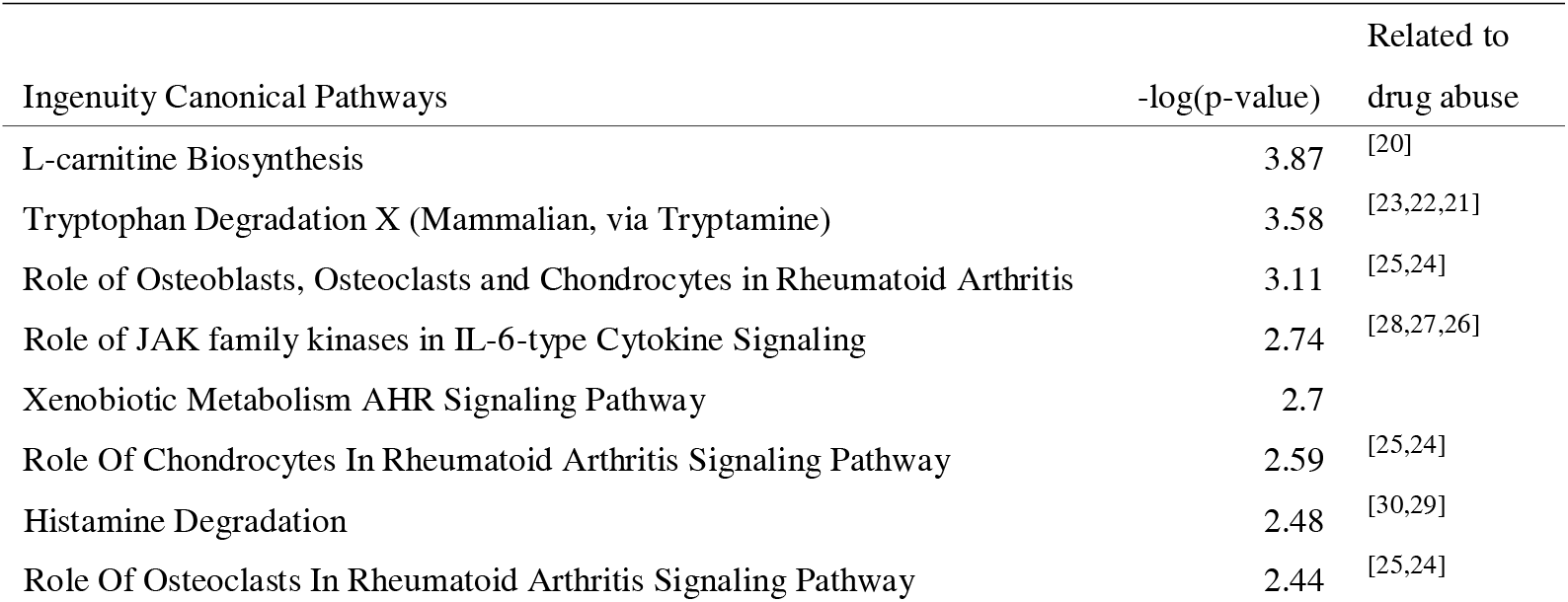
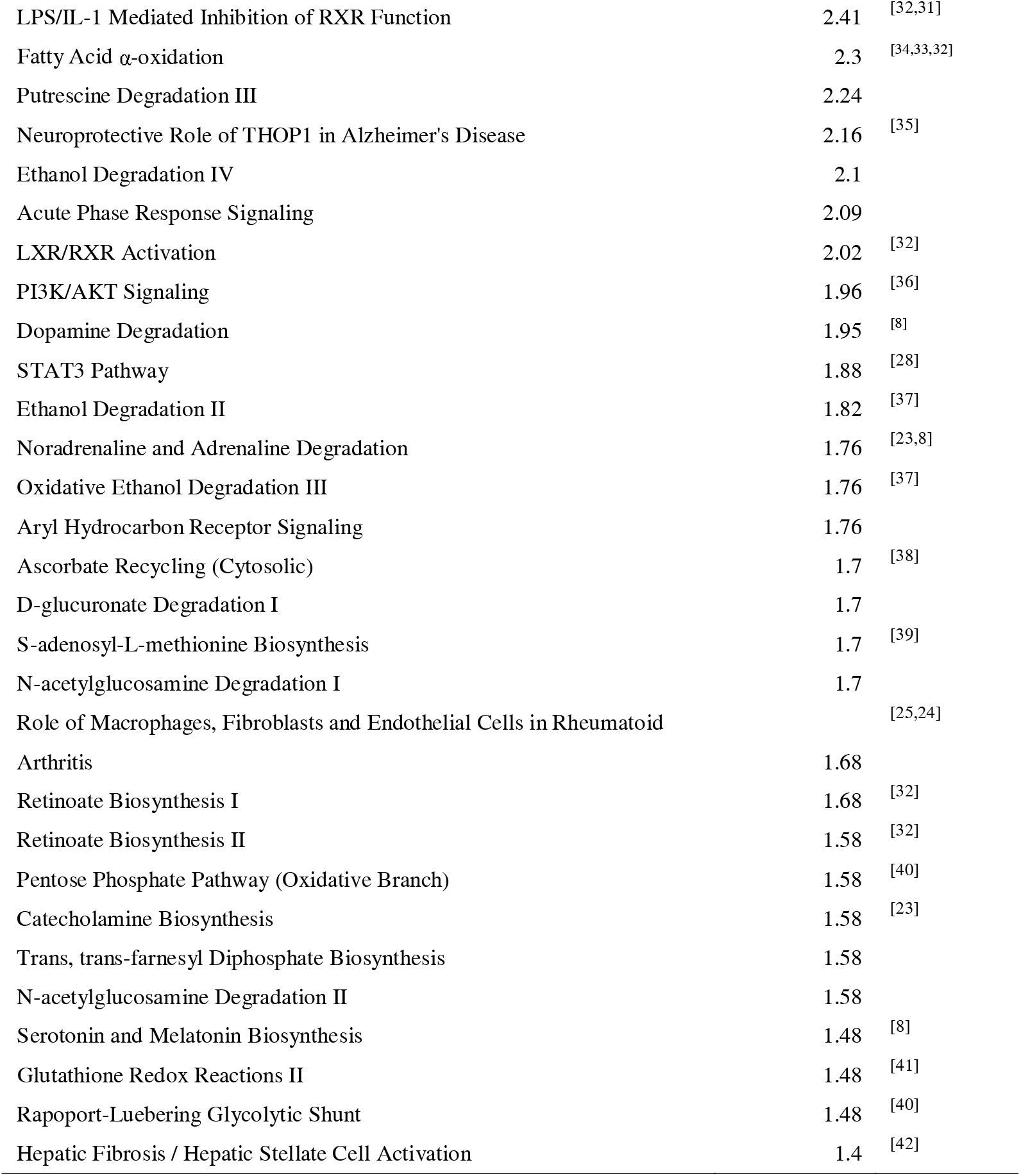
Differential proteins enriched to the IPA pathway between patients in the acute phase and healthy subjects.

The enriched IPA pathways include the metabolism of many important neurotransmitters, their precursors, derivatives, and neuromodulators, such as L-carnitine biosynthesis, tryptophan degradation, histamine degradation, fatty acid alpha-oxidation, putrescine degradation, neuroprotective role of THOP1 in Alzheimer’s disease, dopamine degradation, ethanol degradation, norepinephrine and epinephrine degradation, aryl hydrocarbon receptor signaling transduction, ascorbic acid recycling, S-adenosyl-L-methionine (SAM) biosynthesis, retinoate biosynthesis, catecholamine biosynthesis, serotonin and melatonin biosynthesis, and glutathione redox reactions.

L-carnitine biosynthesis is the pathway with the smallest p-value (greatest significance). L-carnitine is a class of amino acids that prompts the conversion of fat into energy. Long-term use of methamphetamine and opioid drugs can lead to loss of appetite and weight loss, which the author believes may be related to L-carnitine metabolism. L-carnitine may be able to exert its neuroprotective effects against methamphetamine toxicity through actions at the level of dopamine release or mitochondrial functioning[20].

Three differential proteins are enriched to tryptophan degradation. Tryptophan metabolites are amine precursors of the 5-hydroxytryptamine, melatonin, kynurenine, and quinoline pathways, and their levels have been found to be affected by methamphetamine addiction[21]. The kynurenine pathway is the primary pathway for tryptophan degradation, and studies have found that regulating kynurenine metabolism at certain stages can reduce, prevent, or eliminate drug-like seeking behavior[22].

According to the literature, the mechanism of action of methamphetamine is increased release of catecholamine neurotransmitters such as norepinephrine[23]. Dopamine and serotonin are neurotransmitters that play an important role in the addiction process. Addictive drugs trigger changes in histamine levels within the hypothalamus, striatum, and other parts of the body, and histamine has an important role in the drug addiction process[30,29] Histamine plays an important role in the process of drug addiction.

Four differential proteins were enriched to the neuroprotective role of THOP1 in Alzheimer’s disease with a z-score of −1. THOP1 (thimet oligopeptidase 1) is an enzyme whose main function is the degradation of neuropeptides and hormones, such as oxytocin and angiotensin II, etc. THOP1 plays an important role in the neuromodulation of the central nervous system and the regulation of the immune system, and is closely related to the development and prognosis of many diseases such as Alzheimer’s disease. Methamphetamine abuse may lead to premature development of Alzheimer’s disease and neurodegeneration[35].

L-ascorbic acid prevents methamphetamine-induced neurotoxicity in cortical cells by inhibiting oxidative stress, autophagy and apoptosis[38]. Cocaine significantly reduced S-adenosyl-L-methionine (SAM)/S-adenosyl-L-homocysteine (SAH) ratio levels in the dorsal striatum[39]. Chronic methamphetamine self-administration increased midbrain limbic mitochondrial oxygen consumption and decreased striatal glutathione[41].

Enrichment of several proteins to osteoblasts, osteoclasts, and chondrocytes in rheumatoid arthritis. Methamphetamine causes an increase in dopamine in the central nervous system. The dopaminergic system strongly influences the progression of rheumatoid arthritis[24]. Dopamine may be involved in bone formation, bone remodeling in rheumatoid arthritis, and joint erosion[25].

The enriched IPA pathways also contained some signaling pathways, such as, the role of JAK family kinases in IL-6 type cytokine signaling, xenobiotic metabolic AHR signaling pathway, LPS/IL-1-mediated inhibition of RXR function, acute-phase response signaling, LXR/RXR activation, PI3K / AKT signaling, and STAT3 pathway.

Four differential proteins were enriched to the role of JAK family kinases in IL-6-type cytokine signaling with a z-score of −2. In addition, four differential proteins were enriched to the STAT3 pathway. Serum TNF-α, IL-6 and IL-18 levels were elevated in chronic methamphetamine users[27,26]. It has also been shown that TNF-α and IL-6 have a protective effect on methamphetamine-induced microglial cell death through the IL-6 receptor, particularly through activation of the JAK-STAT3 pathway, which alters pro- and anti-apoptotic proteins[28].

Four differential proteins are enriched to heterologous metabolic AHR signaling pathway and aryl hydrocarbon receptor signaling. The aryl hydrocarbon receptor (AHR) recognizes a number of xenobiotics and natural compounds, such as tryptophan metabolites.

Six differential proteins enriched to LPS/IL-1-mediated inhibition of RXR function. It was shown that protein levels of IL-1β were significantly increased in the hippocampal tissue of methamphetamine-exposed mice[31]. Retinoid X receptor (RXR) is involved in amphetamine-induced locomotor activity[32].

Five differential proteins enriched for PI3K/AKT signaling. Methamphetamine can mediate neuroprotection by activating the dopamine/PI3K/AKT signaling pathway[36].

Four differential proteins were enriched to hepatic fibrosis/hepatic stellate cell activation. Serum levels of alanine aminotransferase and aspartate aminotransferase activities were significantly elevated in rats injected intraperitoneally with methamphetamine compared with controls, suggesting significant liver injury[42].

Glycolysis-related pathways, and fatty acid oxidation were also enriched to. Methamphetamine causes altered carbohydrate metabolism[40]. Methamphetamine inhibits glucose uptake in human neurons and astrocytes. Methamphetamine inhibits glucose uptake by human neurons and astrocytes, and during glucose limitation these cells adapt to fatty acid oxidation as an alternative energy source[33]. Methamphetamine inhibits glucose uptake in human neurons and astrocytes. Methamphetamine induces structural changes in astrocytes through multiple targets, of which cytoarchitecture, steroid biosynthesis, and fatty acid biosynthesis may play important roles in neurological injury[43]. Amphetamine exerts fat mobilization through endogenous catecholamine release[34].

#### 3.2.3 GO analysis and KEGG pathway analysis of differential proteins

Functional analysis of differential proteins using the DAVID database enriched a total of 26 biological processes (BPs) including regulation of gene silencing, regulation of gene expression, epigenetics, telomere organization, DNA replication-dependent nucleosome assembly, nucleosome assembly, negative regulation of endopeptidase activity, cellular adhesion, regulation of presynaptic assembly, bone mineralization, prostaglandin metabolic processes, synapse assembly positive regulation, negative regulation of neuronal death, positive regulation of protein metabolic processes, positive regulation of cell-substrate adhesion, memory, carnitine biosynthetic processes, developmental processes involving reproduction, osteoclast proliferation, protein hydrolysis, viral entry into host cells, angiogenesis, regulation of protein localization to membranes, neural crest cell migration.

Ten differential proteins are enriched to biological processes such as regulation of gene silencing, regulation of gene expression (epigenetic), telomere organization, DNA replication-dependent nucleosome assembly, nucleosome assembly, etc. DNA methylation may be a key molecular mechanism for the long-term persistence of addictive behaviors, primarily through the addition of methyl groups to cytosines in the promoter regions of genes, which impede the binding of transcription factors and thus inhibit gene expression over long time scales[44].

Three differential proteins enriched to prostaglandin metabolic processes. Studies have shown that prostaglandins are drugs that modulate brain disease processes in a positive or negative manner. Prostaglandins (PGs) are formed by sequential oxidation of arachidonic acid under physiological and pathological conditions. For the production of PGs, cyclooxygenase is an essential enzyme with two isoforms named cyclooxygenase-1 (COX-1) and cyclooxygenase-2 (COX-2)[41]. Cyclooxygenase-2 is essential for methamphetamine-induced neurotoxicity[45]. Cannabis patients have been found to possess higher levels of prostaglandin F1a in their serum metabolome than controls[46].

Six differential proteins enriched to angiogenesis. Methamphetamine administration induces retinal hypoxia and angiogenesis[47].

Several differential proteins are enriched to the regulation of synapses, regulation of neuronal death, memory, neural crest cell migration, and other biological processes.

**Table 3.** Biological processes enriched to differential proteins between patients in the acute phase and healthy individuals (BP)

| Term | Count | % | P-Value |
| --- | --- | --- | --- |
| regulation of gene silencing | 10 | 6.6 | 3E-18 |
| regulation of gene expression, epigenetic | 10 | 6.6 | 1.5E-13 |
| telomere organization | 10 | 6.6 | 2.3E-13 |
| DNA replication-dependent nucleosome assembly | 10 | 6.6 | 1.3E-12 |
| nucleosome assembly | 10 | 6.6 | 2.2E-06 |
| negative regulation of endopeptidase activity | 7 | 4.6 | 0.00079 |
| cell adhesion | 13 | 8.6 | 0.00093 |
| regulation of presynapse assembly | 4 | 2.6 | 0.0024 |
| heterophilic cell-cell adhesion via plasma membrane cell adhesion molecules | 4 | 2.6 | 0.0069 |
| bone mineralization | 4 | 2.6 | 0.0076 |
| prostaglandin metabolic process | 3 | 2 | 0.011 |
| positive regulation of synapse assembly | 4 | 2.6 | 0.011 |
| negative regulation of neuron death | 4 | 2.6 | 0.012 |
| positive regulation of protein metabolic process | 3 | 2 | 0.014 |
| positive regulation of cell-matrix adhesion | 3 | 2 | 0.018 |
| memory | 4 | 2.6 | 0.028 |
| carnitine biosynthetic process | 2 | 1.3 | 0.029 |
| cell-substrate junction assembly | 2 | 1.3 | 0.037 |
| developmental process involved in reproduction | 2 | 1.3 | 0.037 |
| osteoclast proliferation | 2 | 1.3 | 0.037 |
| proteolysis | 8 | 5.3 | 0.037 |
| Homophilic cell adhesion via plasma membrane adhesion molecules | 5 | 3.3 | 0.04 |
| viral entry into host cell | 4 | 2.6 | 0.041 |
| angiogenesis | 6 | 3.9 | 0.041 |
| regulation of protein localization to membrane | 2 | 1.3 | 0.044 |
| neural crest cell migration | 3 | 2 | 0.048 |

Differential proteins were enriched to 22 molecular functions, including calcineurin binding, structural components of chromatin, protein heterodimerization activity, protein binding, serine-type endopeptidase inhibitor activity, interleukin-1 receptor activity, amino butyric acid dehydrogenase activity, metalloamylopeptidase activity, aminopeptidase activity, viral receptor activity, fatty acid binding, muscle structural components, peptidase activity, and actin binding, Glyceraldehyde-3-phosphate dehydrogenase (NAD +) (non-phosphorylated) activity, serine-type peptidase activity, dipeptidyl peptidase activity, neuregulin binding, catalytic activity, myosin binding, oxidoreductase activity.

Differential proteins were enriched to 10 KEGG pathways including Shigellosis, systemic lupus erythematosus, alcoholism, transcriptional dysregulation in cancer, neutrophil extracellular trap formation, metabolic pathways, beta-alanine metabolism, osteoclast differentiation, cell adhesion molecules, and cytokine-cytokine receptor interactions.

11 differential proteins enriched to alcoholism. Studies have shown similarities in the effects of drugs of abuse such as alcohol and psychoactive substances, such as reduced dopaminergic markers[37].

Three differential proteins enriched to β-alanine metabolize this pathway. β-alanine is a neuromodulatory substance. four cyclohexene analogs of γ-aminobutyric acid (GABA) and β-alanine have been engineered to serve as conformationally rigid analogs of the epileptic and drug-addictive drug aminocaproic acid, and to serve as potential mechanism-based inactivators of the enzyme γ-aminobutyric acid transaminase (GABA-AT)[48].

Four differential proteins were enriched to BP such as bone mineralization and osteoclast proliferation. five differential proteins were enriched to osteoclast differentiation, a KEGG pathway. Studies have shown that in a high percentage of methamphetamine abusers, there is a considerable loss of bone mineralization[49].

7 differential protein enrichment to cytokine-cytokine receptor interactions. Changes in cytokine homeostasis are associated with damage to the blood-brain barrier, leading to altered brain plasticity and lasting neurotoxicity[76].

**Table 4.** KEGG pathways enriched to differential proteins between patients in the acute phase and healthy subjects.

| Term | Count | % | P-Value |
| --- | --- | --- | --- |
| Shigellosis | 13 | 8.6 | 5.80E-06 |
| Systemic lupus erythematosus | 10 | 6.6 | 8.50E-06 |
| Alcoholism | 11 | 7.2 | 1.70E-05 |
| Transcriptional misregulation in cancer | 11 | 7.2 | 2.10E-05 |
| Neutrophil extracellular trap formation | 10 | 6.6 | 1.20E-04 |
| Metabolic pathways | 26 | 17.1 | 7.70E-03 |
| beta-Alanine metabolism | 3 | 2 | 3.80E-02 |
| Osteoclast differentiation | 5 | 3.3 | 4.00E-02 |
| Cell adhesion molecules | 5 | 3.3 | 7.30E-02 |
| Cytokine-cytokine receptor interaction | 7 | 4.6 | 7.80E-02 |

### 3.3 Comparative analysis between the rehabilitation group and the healthy group

#### 3.3.1 IPA pathway analysis of differential proteins

In the comparative analysis between the healthy group and the group of recovering patients, 312 differential proteins were screened for FC >1.5 or <0.67, P < 0.05.

Differential protein enrichment to 62 IPA pathways.

The pathways with the greatest significance were FXR / RXR activation, LXR / RXR activation. The primary substrate for amphetamine (AMPH)-induced motor activity is associated with dopamine forebrain circuits. Brain regions associated with AMPH-induced motor activity express high levels of retinoid receptors (RXR), and retinoid X receptors are involved in amphetamine-induced motor activity[32].

Synaptogenesis signaling pathways, axon guidance signals are enriched to. Methamphetamine causes disruption of dopaminergic nerve endings in the mammalian forebrain, affecting neuronal growth and synapse formation[8,75].

Differential proteins are also enriched for many immune-related pathways, such as coagulation, lymphocyte signaling, complement system, rheumatoid arthritis signaling, and systemic lupus erythematosus signaling.

Six differential proteins were enriched for coagulation with a z-score of 0.8. The plasminogen activation pathway was also enriched. Opioid addiction leads to alterations in the coagulation system[50]. 4 differential proteins were enriched to the complement system with a z-score of −1. The complement system is involved in methamphetamine addiction[51], synthetic psychoactive drug abuse may trigger multiple complement system mutations[52]. The complement system is involved in methamphetamine addiction.

Three differential proteins are enriched for sucrose degradation. Sucrose ingestion activates the mesocortical limbic system in a manner similar to substance abuse[53]. Several differential proteins are enriched to glycolysis, gluconeogenesis-related pathways. four differential proteins are enriched to juvenile-onset adult-onset diabetes/adult-onset diabetes of the young (MODY) signaling. six differential proteins are enriched to iron-homeostasis signaling pathways, and five differential proteins are enriched to iron-death signaling pathways. Methamphetamine causes a toxic syndrome characterized by altered carbohydrate metabolism, dysregulation of calcium and iron homeostasis, increased oxidative stress, and disruption of mitochondrial function[40].

Differential proteins are enriched to a number of important amino acid biosynthesis or degradation processes, including alanine biosynthesis, alanine degradation, glycine biosynthesis, tryptophan degradation, methionine recycling, and tyrosine synthesis. The metabolic profile of these amino acids may be relevant to the addiction process.

Glycine transporter protein (GlyT)-1 plays a key role in maintaining glycine levels at glutamatergic synapses.Glycine is a variant agonist of the N-methyl-D-aspartate (NMDA) receptor, and activation of the NMDA receptor is an important step in the induction of methamphetamine dependence and psychosis[54]. Glycine reduces methamphetamine-induced motor activity[55].

Significant changes in alanine, aspartate, and glutamate metabolic pathways, and cysteine and methionine metabolic pathways in the plasma metabolome of methamphetamine abusers withdrawing from the program[56].

Dopaminergic neurons are deeply involved in addiction, with tyrosine hydroxylase catalyzing the first and rate-limiting step in dopamine (DA) biosynthesis. Overexpression of tyrosine hydroxylase in dopaminergic neurons increases sensitivity to methamphetamine. Tyrosine hydroxylase may be a pharmacological target for improving sensitivity to drugs of abuse[57].

Anandamide degradation of this pathway was enriched to. anandamide, also known as arachidonicotinoylethanolamide, was the first endogenous cannabinoid to be discovered. Substance abuse leads to disruption of synaptic plasticity in the brain circuits involved in addiction, with alterations in normal endogenous cannabinoid activity playing an important role. This promotes the development of abnormal brain changes and addictive behaviors characteristic of substance use disorders[58].. Several studies have shown that Anandamide has an overall modulatory effect on brain reward circuits. Some reports suggest that it is involved in addictive behaviors of other drugs of abuse and can also act as a behavioral reinforcer in animal models of drug abuse. All of these effects of Anandamide appear to be enhanced by pharmacological inhibition of its metabolic degradation, and treatment with fatty acid amide hydrolase inhibitors, the primary enzyme responsible for its degradation, resulted in elevated brain levels of Anandamide that appears to affect the rewarding and reinforcing effects of many drugs of abuse[59].

Many signaling-related pathways were enriched to.

16 differential proteins enriched for p70S6K signaling. Mitogen-stimulated p70-S6 kinase (p70-S6K) is involved in the development of sensitization to methamphetamine-induced reward effects in rats[60]. p70S6K is now recognized as a novel therapeutic target for drug development in diseases such as cancer.

Twelve differential proteins were enriched for IL-15 signaling, and seven differential proteins were enriched for IL-15 production. Methamphetamine (METH) addiction and withdrawal cause severe damage to the immune and nervous systems, and patients in acute withdrawal from methamphetamine had significantly lower levels of interleukin (IL)-1β, IL-9, and IL-15[61].

5 differential proteins enriched to Ephrin B signaling with a z-score of −1.3. Ephrin is an axon guidance molecule that may be associated with opioid addiction-related behaviors[62]. the Eph receptor A4 plays a role in demyelination and depression-related behaviors[63].

10 differential proteins enriched to grid protein (Clathrin)-mediated endocytosis signaling. Endocytosis of dopamine receptors is regulated by many components, such as lattice proteins, β-inhibitory proteins, fiducial proteins, and Rab family proteins. Dopamine receptors escape from lysosomal digestion and their recycling occurs rapidly, enhancing dopaminergic signaling[64].

Eight differential proteins enriched to the actin cytoskeleton signal with a z-score of −0.4. literature suggests that the actin cytoskeleton may serve as a therapeutic target for methamphetamine relapse prevention[65]. Methamphetamine decreases expression of tight junction proteins, rearranges the F-actin cytoskeleton, and increases blood-brain barrier permeability through a RhoA/ROCK-dependent pathway[66].

Nine differential proteins enriched to sperm motility. Methamphetamine affects intracellular calcium homeostasis by acting through calcium signaling pathway-related proteins. In addition, it may disrupt ionic homeostasis in spermatozoa through GABA A-α1 receptors and calcium-binding proteins, triggering changes in intracellular calcium and chloride ions that are associated with sperm motility[67].

Five differential proteins were enriched for Macropinocytosis signaling. Overstimulation of macropinocytosis during methamphetamine exposure in SH-SY5Y human neuroblastoma cells resulted in lysosomal dysfunction[68].

17 differential proteins enriched for phospholipase C signaling. Studies have shown that the central renin-angiotensin system is associated with neurological disorders. Inhibition of PLCβ1 effectively mitigated methamphetamine-induced neurotoxicity and methamphetamine self-administration, and the reinforcing and motivating effects of methamphetamine were significantly reduced by central blockade of PLCβ1 involved signaling pathway[69].

6 differential proteins enriched to Th1, Th2 activation pathways. Cocaine use disorders are associated with changes in Th1/Th2/Th17 cytokines and lymphocyte subsets[70].

Seven differential proteins enriched for RHOGDI signaling. Rho GDP-dissociation inhibitor (Rho GDI) identified as a down-regulated regulator of Rho family GTPases[71]. Rho kinase inhibitors ameliorate cognitive deficits in a methamphetamine-induced male mouse model of schizophrenia[72].

Role of 14 differential proteins enriched to nuclear factor of activated T cells (NFAT) in the regulation of immune responses. Methamphetamine induces calmodulin phosphatase activation, nuclear translocation of NFAT[73], upregulation of the calmodulin/NFAT-induced Fas ligand/Fas death pathway involved in methamphetamine-induced neuronal apoptosis[74] The up-regulation of Fas ligand/Fas death pathway induced by calmodulin/NFAT is involved in methamphetamine-induced apoptosis.

In addition, in the comparative analysis of patients in the recovery phase and healthy subjects, a number of pathways were found to overlap with those enriched in the comparison of patients in the acute phase and healthy subjects. The overlapping pathways included LXR/RXR activation, acute phase response signaling, neuroprotective role of THOP1 in Alzheimer’s disease, the role of the JAK family of kinases in IL-6-type cytokine signaling, hepatic fibrosis/hepatic stellate cell activation, role of chondrocytes in rheumatoid arthritis signaling pathways, PI3K/AKT signaling, STAT3 pathway, tryptophan degradation, L-carnitine biosynthesis, LPS/IL-1-mediated inhibition of RXR function, and role of macrophages, fibroblasts, and endothelial cells in rheumatoid arthritis. As shown in the Table 5, "-" indicates that there are no duplications and are pathways unique to recovering patients. Many of these pathways may have relevance to methamphetamine, addiction.

**Table 5.** Differential proteins enriched to the IPA pathway between recovered patients and healthy subjects (RH indicates recovered patients compared with healthy subjects, AH indicates acute patients compared with healthy subjects)

| Ingenuity Canonical Pathways | RH -log(p-value) | AH -log(p-value) | References |
| --- | --- | --- | --- |
| FXR/RXR Activation | 12.4 | - | [32] |
| LXR/RXR Activation | 11.4 | 2.02 | [32] |
| Acute Phase Response Signaling | 11.3 | 2.09 |  |
| Neuroprotective Role of THOP1 in Alzheimer's Disease | 5.17 | 2.16 | [35] |
| Coagulation System | 4.93 | - | [50] |
| p70S6K Signaling | 4.59 | - | [60] |
| PI3K Signaling in B Lymphocytes | 3.94 | - | [36] |
| B Cell Development | 3.9 | - |  |
| Role of JAK family kinases in IL-6-type Cytokine Signaling | 3.83 | 2.74 | [28,27,26] |
| Sucrose Degradation V (Mammalian) | 3.76 | - | [53] |
| IL-15 Signaling | 3.74 | - | [61] |
| FcγRIIB Signaling in B Lymphocytes | 3.63 | - |  |
| IL-15 Production | 3.36 | - | [61] |
| Hepatic Fibrosis / Hepatic Stellate Cell Activation | 3.24 | 1.4 | [42] |
| Ephrin Receptor Signaling | 3.05 | - | [63,62] |
| Clathrin-mediated Endocytosis Signaling | 2.97 | - | [64] |
| Systemic Lupus Erythematosus In B Cell Signaling Pathway | 2.9 | - |  |
| Complement System | 2.71 | - | [52,51] |
| Synaptogenesis Signaling Pathway | 2.62 | - | [8,75] |
| Axonal Guidance Signaling | 2.52 | - | [8,75] |
| Intrinsic Prothrombin Activation Pathway | 2.5 | - | [50] |
| B Cell Receptor Signaling | 2.44 | - |  |
| Ephrin B Signaling | 2.36 | - | [63,62] |
| Role Of Chondrocytes In Rheumatoid Arthritis Signaling Pathway | 2.3 | 2.59 | [25,24] |
| Pathogen Induced Cytokine Storm Signaling Pathway | 2.29 | - |  |
| Glycolysis I | 2.28 | - | [40] |
| Macropinocytosis Signaling | 2.26 | - | [68] |
| Gluconeogenesis I | 2.23 | - | [40] |
| SPINK1 Pancreatic Cancer Pathway | 2.04 | - |  |
| Glycine Betaine Degradation | 2.04 | - |  |
| PI3K/AKT Signaling | 2.01 | 1.96 | [36] |
| Sperm Motility | 1.94 | - | [67] |
| Iron homeostasis signaling pathway | 1.85 | - | [40] |
| STAT3 Pathway | 1.8 | 1.88 | [28] |
| Phospholipase C Signaling | 1.75 | - | [69] |
| Communication between Innate and Adaptive Immune Cells | 1.65 | - |  |
| Extrinsic Prothrombin Activation Pathway | 1.64 | - | [50] |
| Osteoarthritis Pathway | 1.63 | - | [25,24] |
| Altered T Cell and B Cell Signaling in Rheumatoid Arthritis | 1.6 | - | [25,24] |
| Actin Cytoskeleton Signaling | 1.56 | - | [66,65] |
| VDR/RXR Activation | 1.55 | - | [32] |
| Maturity Onset Diabetes of Young (MODY) Signaling | 1.55 | - | [40] |
| Alanine Degradation III | 1.53 | - | [56] |
| Alanine Biosynthesis II | 1.53 | - | [56] |
| Glycine Biosynthesis I | 1.53 | - | [55,54] |
| Tryptophan Degradation X (Mammalian, via Tryptamine) | 1.5 | 3.58 | [23,22,21] |
| Systemic Lupus Erythematosus Signaling | 1.49 | - |  |
| Th1 and Th2 Activation Pathway | 1.41 | - | [70] |
| Ferroptosis Signaling Pathway | 1.38 | - | [40] |
| IL-6 Signaling | 1.36 | - | [28,27,26] |
| L-carnitine Biosynthesis | 1.36 | 3.87 | [20] |
| Guanine and Guanosine Salvage I | 1.36 | - |  |
| Methionine Salvage II (Mammalian) | 1.36 | - | [56] |
| Anandamide Degradation | 1.36 | - | [59,58] |
| Tyrosine Biosynthesis IV | 1.36 | - | [57] |
| RHO GDI Signaling | 1.35 | - | [72,71] |
| Hepatic Fibrosis Signaling Pathway | 1.34 | - | [42] |
| Granulocyte Adhesion and Diapedesis | 1.34 | - |  |
| Role of NFAT in Regulation of the Immune Response | 1.33 | - | [74,73] |
| LPS/IL-1 Mediated Inhibition of RXR Function | 1.32 | 2.41 | [32,31] |
| Th2 Pathway | 1.31 | - | [70] |
| Role of Macrophages, Fibroblasts and Endothelial Cells in Rheumatoid Arthritis | 1.31 | 1.68 | [25,24] |

THOP1 (thimet oligopeptidase 1) is an enzyme whose main function is the degradation of neuropeptides and hormones such as oxytocin and angiotensin II. THOP1 plays an important role in the neuromodulation of the central nervous system and the regulation of the immune system, and is closely related to the development and prognosis of many diseases such as Alzheimer’s disease. Methamphetamine abuse may contribute to the premature development of Alzheimer’s disease and neurodegeneration[35].

Elevated serum TNF-alpha, IL-6 and IL-18 levels in chronic methamphetamine users[27,26]. It has also been shown that TNF-α and IL-6 have a protective effect on methamphetamine-induced microglial cell death through the IL-6 receptor, particularly through activation of the JAK-STAT3 pathway, which alters pro- and anti-apoptotic proteins[28].

Methamphetamine can mediate neuroprotection by activating the dopamine/PI3K/AKT signaling pathway[36]. L-carnitine may be able to exert its neuroprotective effects against methamphetamine toxicity by acting at the level of dopamine release or mitochondrial functioning[20].

Levels of tryptophan metabolites, which are amine precursors of the 5-hydroxytryptamine, melatonin, kynurenine, and quinoline pathways, have been found to be affected by methamphetamine addiction[21]. The kynurenine pathway is the primary pathway for tryptophan degradation, and studies have found that regulating kynurenine metabolism at certain stages can reduce, prevent, or eliminate drug-like drug-seeking behaviors[22].

Methamphetamine addiction is strongly associated with the dopaminergic system. The dopaminergic system strongly influences the progression of rheumatoid arthritis[24]. Dopamine may be involved in bone formation, bone remodeling in rheumatoid arthritis, and joint erosion[25].

We found that some of the differential proteins and pathways in people who had been abstinent for more than three months (comparisons between recovering patients and healthy individuals) overlapped with those of people who were currently using drugs (comparisons between acute-phase patients and healthy individuals), and these consistencies may reflect the fact that, even after more than three months of abstinence from methamphetamine, patients continue to be left with long-lasting effects in their bodies that do not return to the level of a healthy person, and that these effects may also explain the continued high rate of relapse in patients who have quit using methamphetamine. These effects may also explain why relapse rates remain high among patients after methamphetamine withdrawal. However, it is also possible for methamphetamine withdrawal patients to return to healthy levels after a longer period of abstinence. In addition to the overlapping pathways, many other pathways likewise show relevance to addiction. These differential proteins and pathways may be able to provide potential drug targets for the treatment of drug addiction and provide clues for the exploration of the mechanisms of drug addiction, and we look forward to further exploration in subsequent studies.

#### 3.3.2 GO analysis and KEGG pathway analysis of differential proteins

Differential proteins were enriched to 83 biological processes (p < 0.05), including negative regulation of endopeptidase activity, cell adhesion, axon guidance, multicellular biogenesis, positive regulation of cell migration, positive regulation of kinase activity, positive regulation of synapse assembly, carbohydrate metabolic processes, protein hydrolysis, Ephrin receptor signaling pathway, complement activation, classical pathway, intercellular adhesion, receptor mediated endocytosis, transmembrane receptor protein tyrosine kinase signaling pathway, innate immune response, negative chemotaxis, peptidyl-tyrosine phosphorylation, coagulation, angiogenesis, negative regulation of fibrinolysis, immune response, regulation of extracellular matrix disassembly, axon myofascial tremor, glycosidolytic metabolic processes, induction of bacterial agglutination, immunoglobulin production, epithelial cell differentiation, retinal homeostasis, virus entry into the host cell, inflammatory response, neural crest cell migration, negative regulation of fibrinogen activation, cell matrix adhesion, antimicrobial peptide-mediated antimicrobial humoral immune response, fructose metabolism processes, negative regulation of cysteine-type endopeptidase activity, regulation of blood coagulation, nucleotide metabolism processes, fibrinogen activation, adaptive immune response, cytosolic iron homeostasis, glycosaminoglycan catabolism processes, defense response to gram-positive bacteria, thyroid hormones, and other factors. defense response, thyroid hormone transport, defense response against fungi, negative regulation of coagulation, dendritic spine development, transforming growth factor beta receptor signaling pathway, positive regulation of viral entry into the host cell, positive regulation of interleukin-6 production, proteolytic metabolic processes, response to hypoxia, acute phase response, positive regulation of tumor necrosis factor production, phospholipid homeostasis, cellular migration, calcium-dependent Intercellular adhesion via plasma membrane cell adhesion molecules, intercellular signaling, smooth muscle cell-matrix adhesion, glycolipid catabolic processes, iron ion homeostasis, positive regulation of matrix adhesion-dependent cell spreading, retinal ganglion cell axon guidance, cytokine-mediated signaling pathways, iron ion transport, neutrophil chemotaxis, positive regulation of phagocytosis, positive regulation of angiogenesis, complement dependent cytotoxicity, cytoplasmic actin-based contraction involved in cell motility, zymogen activation, myelin sheath formation, lysosomal organization, etc. The results of the IPA pathway analysis partially overlap with those of the GO analysis using the DAVID database, and are able to verify and complement each other.

The biological processes that differential proteins are enriched for include many neuromodulation-related biological processes, such as axon guidance, synapse assembly, neural spine cell migration, neuronal development, dendritic spine development, and myelin sheath formation. It also includes many biological processes related to immunity, such as complement activation, innate immune response, immune response, immunoglobulin production, and inflammatory response. Methamphetamine has the ability to modulate immune cells[76], which has significant effects on the brain, immunity, and digestion[77]. The effects of methamphetamine on the brain, immunity, and digestion are significant.

In the comparative analysis of patients in the recovery phase with healthy subjects, a number of biological processes were found to overlap with those enriched in the comparison of patients in the acute phase with healthy subjects, and the overlapping biological processes included negative regulation of endopeptidase activity, cell adhesion, positive regulation of synaptic assembly, protein hydrolysis, heterophilic cell-to-cell adhesion via plasma membrane cell adhesion molecules, homophilic cell adhesion via plasma membrane adhesion molecules, angiogenesis, viral entry into host cells, and neural crest cell migration. Many biological processes may be relevant to addiction.

Seventeen differential proteins are enriched for the regulation of endopeptidase activity. Many endopeptidases regulate the degradation of neuropeptides, enkephalins, which are associated with the regulation of mood, anxiety, reward, euphoria, and pain[78]. Among the differential proteins, carboxypeptidase E (CPE) (FC=0.55,P=0.02) regulates dopamine transporter protein activity[79].

13 differential proteins were enriched to angiogenesis. Biological processes such as retinal homeostasis and retinal ganglion cell axon guidance were also enriched to. Methamphetamine administration induced retinal hypoxia and angiogenesis[47].

Seven differential proteins were enriched to the Ephrin receptor signaling pathway. Many axon guidance molecules, such as integrins, semaphorins, and Ephrin, may facilitate oxycodone-induced neuroadaptation by altering axon-target connectivity and synaptogenesis, which may be related to behaviors associated with opioid addiction[62]. the Eph receptor A4 plays a role in demyelination and depression-related behaviors[63]. Eph receptor A4 plays a role in demyelination and depression-related behaviors.

10 differential proteins enriched to the transmembrane receptor protein tyrosine kinase signaling pathway. Receptor tyrosine kinases (RTK) are a large class of proteins that can modulate behavior in the nervous system by regulating neuronal and glial function, and as such, they have been implicated in neurodegenerative diseases and psychiatric disorders such as depression and addiction. Multiple receptor tyrosine kinases regulate alcohol consumption and other behaviors associated with alcohol addiction. Receptor tyrosine kinases are drug targets for the development of treatments for alcohol use disorder (AUD)[80].

**Table 6.** Biological processes enriched to differential proteins between convalescent patients and healthy individuals (BP)

| Term | Count | % | P-Value |
| --- | --- | --- | --- |
| negative regulation of endopeptidase activity | 17 | 5.4 | 1.40E-09 |
| cell adhesion | 31 | 9.9 | 5.10E-09 |
| axon guidance | 16 | 5.1 | 2.40E-07 |
| multicellular organism development | 17 | 5.4 | 3.90E-07 |
| positive regulation of cell migration | 18 | 5.8 | 1.10E-06 |
| positive regulation of kinase activity | 10 | 3.2 | 2.20E-06 |
| positive regulation of synapse assembly | 8 | 2.6 | 5.40E-05 |
| carbohydrate metabolic process | 12 | 3.8 | 9.60E-05 |
| proteolysis | 19 | 6.1 | 1.30E-04 |
| ephrin receptor signaling pathway | 7 | 2.2 | 1.50E-04 |
| complement activation, classical pathway | 10 | 3.2 | 1.60E-04 |
| heterophilic cell-cell adhesion via plasma membrane cell adhesion molecules | 7 | 2.2 | 1.70E-04 |
| receptor-mediated endocytosis | 8 | 2.6 | 2.30E-04 |
| transmembrane receptor protein tyrosine kinase signaling pathway | 10 | 3.2 | 2.70E-04 |
| innate immune response | 23 | 7.4 | 2.70E-04 |
| negative chemotaxis | 6 | 1.9 | 3.60E-04 |
| Homophilic cell adhesion via plasma membrane adhesion molecules | 11 | 3.5 | 4.30E-04 |
| peptidyl-tyrosine phosphorylation | 10 | 3.2 | 4.60E-04 |
| blood coagulation | 8 | 2.6 | 4.90E-04 |
| angiogenesis | 13 | 4.2 | 7.40E-04 |
| positive regulation of peptidyl-tyrosine phosphorylation | 8 | 2.6 | 7.80E-04 |
| negative regulation of fibrinolysis | 4 | 1.3 | 1.00E-03 |
| negative regulation of cell adhesion | 6 | 1.9 | 1.40E-03 |
| immune response | 18 | 5.8 | 2.30E-03 |
| regulation of extracellular matrix disassembly | 3 | 1 | 2.40E-03 |
| axonal fasciculation | 4 | 1.3 | 3.20E-03 |
| fibrinolysis | 4 | 1.3 | 3.20E-03 |
| glycoside catabolic process | 3 | 1 | 3.60E-03 |
| induction of bacterial agglutination | 3 | 1 | 3.60E-03 |
| immunoglobulin production | 7 | 2.2 | 3.80E-03 |
| epithelial cell differentiation | 7 | 2.2 | 3.80E-03 |
| retina homeostasis | 5 | 1.6 | 3.90E-03 |
| viral entry into host cell | 7 | 2.2 | 5.70E-03 |
| inflammatory response | 15 | 4.8 | 6.40E-03 |
| neural crest cell migration | 5 | 1.6 | 6.40E-03 |
| opsonization | 3 | 1 | 6.60E-03 |
| negative regulation of plasminogen activation | 3 | 1 | 6.60E-03 |
| cell-matrix adhesion | 7 | 2.2 | 7.10E-03 |
| Antimicrobial humoral immune response mediated by antimicrobial peptide | 7 | 2.2 | 7.80E-03 |
| ovulation cycle | 3 | 1 | 8.40E-03 |
| fructose metabolic process | 3 | 1 | 1.00E-02 |
| negative regulation of cysteine-type endopeptidase activity | 3 | 1 | 1.00E-02 |
| regulation of blood coagulation | 3 | 1 | 1.20E-02 |
| nucleotide metabolic process | 3 | 1 | 1.20E-02 |
| plasminogen activation | 3 | 1 | 1.20E-02 |
| central nervous system projection neuron axonogenesis | 3 | 1 | 1.20E-02 |
| adaptive immune response | 15 | 4.8 | 1.30E-02 |
| cellular iron ion homeostasis | 5 | 1.6 | 1.30E-02 |
| glycosaminoglycan catabolic process | 3 | 1 | 1.50E-02 |
| Defense response to Gram-positive bacterium | 7 | 2.2 | 1.70E-02 |
| thyroid hormone transport | 3 | 1 | 1.70E-02 |
| Defense response to fungus | 4 | 1.3 | 1.80E-02 |
| negative regulation of blood coagulation | 3 | 1 | 2.00E-02 |
| dendritic spine development | 3 | 1 | 2.00E-02 |
| transforming growth factor beta receptor signaling pathway | 6 | 1.9 | 2.20E-02 |
| positive regulation of viral entry into host cell | 3 | 1 | 2.30E-02 |
| cell-cell junction assembly | 4 | 1.3 | 2.40E-02 |
| positive regulation of interleukin-6 production | 6 | 1.9 | 2.50E-02 |
| protein catabolic process | 5 | 1.6 | 2.50E-02 |
| response to hypoxia | 8 | 2.6 | 2.70E-02 |
| acute-phase response | 4 | 1.3 | 2.90E-02 |
| positive regulation of tumor necrosis factor production | 6 | 1.9 | 2.90E-02 |
| phospholipid homeostasis | 3 | 1 | 2.90E-02 |
| cell migration | 10 | 3.2 | 2.90E-02 |
| calcium-dependent cell-cell adhesion via plasma membrane cell adhesion molecules | 4 | 1.3 | 3.00E-02 |
| cell-cell signaling | 9 | 2.9 | 3.10E-02 |
| smooth muscle cell-matrix adhesion | 2 | 0.6 | 3.10E-02 |
| glycolipid catabolic process | 2 | 0.6 | 3.10E-02 |
| iron ion homeostasis | 4 | 1.3 | 3.20E-02 |
| positive regulation of substrate adhesion-dependent cell spreading | 4 | 1.3 | 3.20E-02 |
| cell-cell adhesion | 8 | 2.6 | 3.30E-02 |
| retinal ganglion cell axon guidance | 3 | 1 | 3.60E-02 |
| cytokine-mediated signaling pathway | 7 | 2.2 | 3.80E-02 |
| iron ion transport | 3 | 1 | 3.90E-02 |
| neutrophil chemotaxis | 5 | 1.6 | 4.10E-02 |
| positive regulation of phagocytosis | 4 | 1.3 | 4.50E-02 |
| positive regulation of angiogenesis | 7 | 2.2 | 4.70E-02 |
| oviduct epithelium development | 2 | 0.6 | 4.70E-02 |
| complement-dependent cytotoxicity | 2 | 0.6 | 4.70E-02 |
| cytoplasmic actin-based contraction involved in cell motility | 2 | 0.6 | 4.70E-02 |
| zymogen activation | 3 | 1 | 4.70E-02 |
| myelination | 4 | 1.3 | 4.70E-02 |
| lysosome organization | 4 | 1.3 | 4.90E-02 |

Differential proteins were enriched to 11 KEGG pathways, including complement and coagulation cascade reactions, pertussis, lysosomes, salivary secretion, axon guidance, hematopoietic cell lines, viruses-herpesviruses, cytokine-cytokine receptor interactions, Staphylococcus aureus infections, glycolysis/glycolysis, and other glycan degradation.

Twelve differential proteins were enriched to complement and coagulation cascade reactions. The complement system is involved in methamphetamine addiction, and complement factor H (CFH) is upregulated in serum from methamphetamine-addicted patients and rats, as well as in certain brain regions in rats[51], synthetic psychoactive drug abuse may trigger multiple complement system mutations[52]. Opioid addiction leads to alterations in the coagulation system[50]. The coagulation system is altered by opioid addiction.

13 differential proteins enriched to cytokine-cytokine receptor interactions. Changes in cytokine homeostasis are associated with damage to the blood-brain barrier, leading to altered brain plasticity and lasting neurotoxicity[76]. In addition, 11 differential proteins were enriched to axon guidance.

**Table 7.** KEGG pathways enriched to differential proteins between convalescent patients and healthy individuals.

| Term | Count | % | P-Value |
| --- | --- | --- | --- |
| Complement and coagulation cascades | 12 | 3.8 | 8.70E-07 |
| Pertussis | 9 | 2.8 | 1.20E-04 |
| Lysosome | 11 | 3.5 | 2.70E-04 |
| Salivary secretion | 8 | 2.5 | 2.40E-03 |
| Axon guidance | 11 | 3.5 | 3.20E-03 |
| Hematopoietic cell lineage | 8 | 2.5 | 3.40E-03 |
| Virion - Herpesvirus | 3 | 0.9 | 1.30E-02 |
| Cytokine-cytokine receptor interaction | 13 | 4.1 | 1.40E-02 |
| *Staphylococcus aureus* infection | 6 | 1.9 | 4.10E-02 |
| Glycolysis / Gluconeogenesis | 5 | 1.6 | 4.30E-02 |
| Other glycan degradation | 3 | 0.9 | 4.80E-02 |

### 3.4 Comparative analysis of the acute and rehabilitation groups

Two hundred differential proteins were screened in the comparative analysis of the acute versus convalescent patient groups, with screening conditions of FC >1.5 or <0.67, P < 0.05.

Functional analysis of differential proteins using the DAVID database enriched a total of 57 biological processes (BPs), including immunoglobulin production, immune response, intercellular adhesion, cell adhesion, noncanonical Wnt signaling pathway, adaptive immune response, synaptic membrane adhesion, cellular response to amino acid stimulation, macromolecular complex assembly, regulation of presynaptic assembly, cell migration, receptor-mediated endocytosis, regulation of cell-substrate adhesion, regulation of cell growth, ossification, positive regulation of myoblast fusion, coagulation, negative regulation of endopeptidase activity, positive regulation of apoptotic processes, catabolic processes of leukotriene D4, substrate adhesion-dependent cell spreading, processes of lipoprotein metabolism, positive regulation of interleukin-6 production, capping of barbed-end actin filaments, localization of proteins to plasma membrane, Protein hydrolysis, extracellular matrix organization, endodermal cell differentiation, regulation of fibrinolysis, TGFβ sequestration in the extracellular matrix, positive regulation of apoptotic processes in keratin-forming cells, development of neuronal projections, negative regulation of cellular proliferation, signal transduction, tissue development, development of the skeletal system, morphogenesis of animal organs, inflammatory response, nervous system development, endocytosis, actin filament polymerization, Acute phase response, multicellular biological development, HDL particle clearance, cytokine-mediated signaling pathways, typical Wnt signaling pathways, positive regulation of phagocytosis, negative regulation of Wnt signaling pathways, antimicrobial peptide-mediated antimicrobial humoral immune response, and regulation of protozoal embryo formation.

Differential proteins were enriched to 27 molecular functions (MFs), including calcium binding, extracellular matrix structural component, signaling receptor activity, NAD + ribonuclease, cyclic ADP ribose production, NAD(P) + ribonuclease activity, carbohydrate binding, enzyme binding, interleukin-1 receptor activity, lipid transport protein activity, antigen binding, Wnt protein binding, NF-κB binding C3b binding, complement component, structural component of the extracellular matrix that confers tensile strength, transmembrane receptor protein tyrosine phosphatase activity, co-receptor activity, biotinidase activity, sequence-specific DNA binding at the proximal region of the core promoter, hormone activity, mannose binding, binding of proteins involved in cell-to-cell adhesion, binding of cellular adhesion molecules, binding of heparin, activity of translation elongation factors, transmembrane signaling receptors, viral receptor activity. activity, viral receptor activity.

Differential proteins were enriched to six KEGG pathways, including complement and coagulation cascade reactions, cytokine-cytokine receptor interactions, cell adhesion molecules, ECM-receptor interactions, hematopoietic cell lines, and amebiasis.

Differential protein enrichment to 56 IPA pathways including liver fibrosis/hepatic stellate cell activation, LXR/RXR activation, B cell development, acute phase response signaling, PI3K signaling in B lymphocytes, pathogen-induced cytokine storm signaling pathway, osteoclasts in rheumatoid arthritis signaling pathway, FXR/RXR activation, IL-15 signaling Role of osteoblasts, osteoclasts and chondrocytes in rheumatoid arthritis, FcγRIIB signaling in B lymphocytes, coagulation system, role of chondrocytes in rheumatoid arthritis signaling pathways, osteoarthritis pathway, systemic lupus erythematosus in the B-cell signaling pathway, wound-healing signaling pathways, clathrin-mediated endocytosis signaling, and the role of osteoclasts in rheumatoid arthritis signaling pathways.) mediated endocytosis signaling, role of macrophages, fibroblasts, and endothelial cells in rheumatoid arthritis, B-cell receptor signaling, IL-10 signaling, Agrin interactions at neuromuscular junctions, neuroprotective role of THOP1 in Alzheimer’s disease, signaling in systemic lupus erythematosus, complement system, and communication between innate and adaptive immune cells, Altered T- and B-cell signaling in rheumatoid arthritis, WNT/β-catenin signaling, JAK family kinases in IL-6-type cytokine signaling, GP6 signaling pathway, leukotriene biosynthesis, NFAT in immune response modulation, idiopathic signaling pathways in pulmonary fibrosis, PPAR signaling, osteoblasts in rheumatoid arthritis signaling pathways. Role of osteoblasts in rheumatoid arthritis signaling pathways, granulocyte adhesion and efflux, myelin signaling pathways, atherosclerosis signaling, IL-6 signaling, VDR/RXR activation, Adult-onset diabetes mellitus of juvenile onset / Adult-onset juvenile diabetes mellitus (MODY) signaling, STAT3 pathway, lung healing signaling pathways, pyrimidine ribonucleotide interconversion, phospholipase C signaling.

### 3.5 Comprehensive Analysis and Discussion

In addition to the group comparative analysis, we also used a one-to-many analysis method, in which each acute-phase patient sample was analyzed in comparison with a group of healthy human samples with essentially corresponding age and sex ratios, to identify differential proteins shared by each acute-phase patient with a view to searching for biomarkers and pathway information related to addiction. Screening conditions were FC > 1.5 or < 0.67, P < 0.05.

The differential protein common to all patients in the acute phase is amphoterin-induced gene and ORF 1 (AMIGO-1), which is expressed in a variety of brain cell types, may regulate dendritic growth and neuronal survival, and may play a role in regeneration and neuroplasticity in the adult nervous system[81]. And neuroplasticity is closely related to the addiction process.

In group comparison of the three groups of samples, AMIGO-1 was also screened as a differential protein, and AMIGO-1 was identified in the three groups as follows: zero in all three groups during the acute phase, a mean of 149 in the rehabilitation group, and a mean of 1,255 in the healthy group. The authors hypothesized that it is possible that the level of AMIGO-1 in the urinary proteome may have shown a negative correlation with methamphetamine use, and that the use of methamphetamine may lead to a decrease in AMIGO-1 content in the urinary proteome, and after withdrawal and rehabilitation, AMIGO-1 content in the urinary proteome increased again. AMIGO-1 may have the potential to be used as a biomarker for methamphetamine use.

It is worth noting that we found that some of the differential proteins and pathways in people who had been abstinent for more than three months (comparisons between recovering patients and healthy individuals) overlapped with those of people who were currently using drugs (comparisons between patients in the acute phase and healthy individuals), and these concordances may reflect the fact that, even after three months of abstinence from methamphetamine or even longer than that, patients continue to be left with persistent effects in the body, which do not return to the healthy individuals’ levels, and these effects may also explain the continued high rate of relapse in patients after methamphetamine withdrawal. In addition to the overlapping pathways, many other pathways also show relevance to addiction. These differential proteins and pathways may be able to provide potential drug targets for the treatment of drug addiction and provide clues for the investigation of the mechanism of drug addiction, and we look forward to further exploration in subsequent studies.

In addition, in the process of analysis, patients in the recovery group were screened to obtain more differential proteins and pathways than patients in the acute group in comparison with healthy people, and the difference was more significant. The reasons for the analysis may be as follows: methamphetamine can have a great impact on the body in a short period of time, physiological indicators will fluctuate drastically over time, the human body’s response under the influence of different doses of drugs, different times of drug use, different metabolic states may be different, and urine is a window, which is very sensitive to the body’s reflection of the change in state in a short period of time. In addition, human susceptibility to addiction varies widely[82]. Therefore, the heterogeneity of samples from patients in the acute phase within 24 hours of acute cessation of methamphetamine use is relatively high. The results of the present study may be beneficial for subsequent refinement of the experimental design by grouping the samples more finely to ensure that there is less variability within groups and that commonalities can be better found.

In conclusion, this study comparatively analyzed the urinary proteomes of methamphetamine-using patients (patients who stopped using methamphetamine for less than 24 hours), methamphetamine-abstinent patients (those who stopped using the drug for more than 3 months and entered rehabilitation treatment), and healthy individuals, and the urine proteomes of methamphetamine-abusing patients were significantly different from those of healthy individuals, with some of the differential proteins and their enrichment into the biological functions were shown to be associated with addiction or methamphetamine neurotoxicity. This study innovatively established an approach to study addictive drugs from the perspective of urine proteomics, demonstrating that the urine proteome can reflect the effects of methamphetamine abuse on the organism in a more systematic and comprehensive manner, and has the potential to provide clues for the research and practice of clinical addictive diseases.

## 4 Perspective

This study is important for understanding the mechanism and biological basis of methamphetamine addiction. By analyzing the changes in the urinary proteome of methamphetamine-addicted patients, we can identify protein biomarkers related to addiction and provide new clues and methods for the early diagnosis and treatment of methamphetamine addiction. In addition, by deeply studying the effects of methamphetamine addiction on the organism, we can reveal some important details of the addiction mechanism, which will help to develop more effective prevention and treatment strategies.

Meanwhile, this study provides a new perspective of urine proteome for the study of addictive substances and offers a proven research method and analysis strategy. With the advantages of high sensitivity, non-invasive collection and stable sample, urine is expected to be one of the important tools for the study of addictive substances.

Despite the significance and potential of this study, there is some room for improvement. Follow-up studies can combine the results of this study with clinical data to further validate that the urinary proteome of methamphetamine-addicted patients has clinical application. Through large-sample clinical studies, we can evaluate the sensitivity, specificity and predictive value of these protein markers and provide reliable biomarkers for the early diagnosis and treatment of methamphetamine addiction.

Finally, the results of this study can also provide guidance for the development of intervention and treatment strategies for methamphetamine addiction. By gaining a deeper understanding of the effects of methamphetamine addiction on the organism, we can target and design interventions to reduce addictive behaviors and alleviate addiction-related physical and psychological problems. Further research can explore the effects of different intervention approaches and look for more effective and individualized treatment options.

In conclusion, this study is of great significance in revealing the mechanism of methamphetamine addiction, searching for biomarkers, and guiding clinical diagnosis and treatment. At the same time, this study provides inspiration for subsequent experimental and clinical studies, and promotes in-depth understanding and effective management of methamphetamine addiction.

## Notes

### Competing Interest Statement

The authors have declared no competing interest.

